# Transgenerational effects induced by thiacloprid in Anterior prostate tissue are associated with alterations in DNA methylation at developmental genes

**DOI:** 10.1101/2025.06.20.660738

**Authors:** Ouzna Dali, Chaima Diba Lahmidi, Tayeb Mohammed Belkhir, Theo De Gestas, Christine Kervarrec, Pierre-Yves Kernanec, Fatima Smagulova

## Abstract

**Background:** Neonicotinoids are widely used pesticides that cause a catastrophic decrease in bee and bumblebee populations worldwide. In addition to insects, neonicotinoids have toxic effects on other species, including lizards, birds, and mammals. Previous studies have shown that gestational exposure to thiacloprid has transgenerational effects on the testis and thyroids.

**Objectives:** In this project, we described the epigenetic effects of thiacloprid on anterior prostate tissue in directly exposed F1 males and nondirectly exposed F3 males

**Methods:** We used paraffin sections for morphological analysis, frozen tissue sections for immunofluorescence analysis, RT‒qPCR for gene expression analysis, histone purification and western blotting for protein analysis, and ChIP‒qPCR for histone H3K4me3 occupancy analysis.

**The results:** We observed a tendency toward an increase in epithelial hyperplasia in F1 and not a significant increase in F3. We detected elevated levels of phosphorylated histone H3 at serine 10, a marker of mitosis, in both the F1 and F3 prostates. A significant increase in the level of the Ki-67 marker of proliferation was detected in the F1 anterior prostate but not in the F3 anterior prostate. *Hox* gene expression was upregulated in F1 anterior prostate and downregulated in F3 anterior prostate. The changes in gene expression were associated with histone H3K4me3 alterations at the promoters of the genes. We determined that regions of *Hox* genes that play important roles in the anterior prostate have altered DNA methylation in the sperm of F1 and F3. These alterations in DNA methylation were negatively related to gene expression.

**Conclusion:** Our analysis revealed that gestational exposure to thiacloprid caused epigenetic alterations in the anterior prostates of F1 males and nondirectly exposed F3 males. DNA methylation changes at the promoters of anterior prostate development genes could be responsible for transgenerational effects in anterior prostates.

## Introduction

Neonicotinoids are widely used pesticides that are intended to control pest populations. The intensive use of neonicotinoids has led to a dramatic decline in bee and bumblebee populations ^1^. In mammals, the toxicity of neonicotinoids has been detected in the central nervous system, specifically concerning the function of acetylcholine receptors ^2^. We and other laboratories have shown that neonicotinoid thiacloprid is toxic for thyroid gland development; it induces alterations in thyroid gland morphology and perturbs the production of thyroid hormones ^3, 4^. We showed that gestational exposure to thiacloprid induces transgenerational alterations in the male reproductive system ^5^. These effects are mediated via changes in DNA methylation at regions known as superenhancers and at germ cell responsive reprogramming genes (GRRs). These GRR genes are essential for the establishment of the germ cell population during embryonic development ^6^. Notably, exposure to thiacloprid reduces the level of testosterone in the blood serum of F3 males ^6^. In this study, we focused on the male accessory gland, the anterior prostate. Previously, neonicotinoids were shown to affect prostate tissue. Specifically, it has been determined that the neonicotinoid imidacloprid (IMI) is toxic to human prostate epithelial cells and induces apoptosis and oxidative stress ^7^. In an *in vivo* study, IMI exposure affected the weight of prostates and led to a decrease in testosterone levels ^8^. The Environmental Protection Agency in the USA (EPA) reported that increased prostate weights and prostatic hypertrophy were induced in dogs after 90 days of treatment with thiacloprid. The EPA classified thiacloprid as “likely to be carcinogenic to humans” based on increased uterine tumors in rats, thyroid follicular adenomas in rats, and ovarian tumors in mice (https://www3.epa.gov/pesticides/chem_search/reg_actions/registration/fs_PC-014019_26-Sep-03.pdf)

Many transcription factors, including a family of homeobox *Hox* genes, play important roles in prostate development. *Hoxa10, Hoxa13, Hoxb13,* and *Hoxd13* have been shown to contribute to the development and regulation of prostate morphogenesis ^9,10,11,12^. The loss of *Hoxb13* function results in impaired epithelial differentiation in the prostate ^11^. Abnormalities in Hoxa10 functional loss lead to altered prostate morphology ^13^. *Nkx3.1* mutations are associated with increased epithelial proliferation in the prostate ^14, 15^.

Prostate gland processes are regulated by epigenetic mechanisms. Deregulation of these genes could lead to pathological processes. Several epigenetic chromatin remodeling factors are overexpressed in prostate cancer, including *Ezh2*^16^, *Hdac1*^17,18^ and *Kdm5a*^19, 20^.

In this study, we aimed to reveal the effects of thiacloprid on the anterior prostate morphology and to gain insight into the epigenetic mechanisms involved in the regulation of gene expression. We showed that gestational exposure to thiacloprid tended to increase epithelial hyperplasia in anterior prostate in F1 but not in F3. We detected increased levels of the mitosis marker PHH3 in both generations, and the expression of the oncogenesis marker Ki-67 was significantly increased in F1 directly exposed males but not in F3. The expression of genes encoding transcription factors, hormones, and chromatin factors was altered in both the F1 and F3 anterior prostates. We observed that gene expression and histone H3K4me3 occupancy at gene promoters consistently changed in similar directions in these mice. Compared with changes in gene expression, alterations in DNA methylation at the promoters of prostate development genes were altered in F1 and F3, mainly in opposite directions.

## Methods

### Mouse treatment and dissections

Outbred Swiss mice (RjOrl) were used for all the experiments. The exposure window was limited to a period between E6.5 and E15.5, which corresponds to the somatic-to-germline transition (SGT) 21. Thiacloprid (Thia) was administered at a dose of 6 mg/kg body weight/day. Thia is distributed throughout the body following exposure and can pass through the placental barrier according to previous studies ^22^. In our study, *thia* was suspended in olive oil and administered via oral gavage at a dose of 6 mg/kg body weight/day in a volume of 150 μl. The control mice were treated with the same volume of oil. For the *control* group, male progeny of F0 females were defined as F1 and were crossed with nonlittermate untreated females to give rise to the F2 generation. Similarly, the male progeny of F2 were crossed with nonlittermate untreated females to give rise to the F3 generation. The oil-treated control animals were processed and crossed in the same way as the treated control animals were. Since we were particularly interested in the transgenerational inheritance of anterior prostate pathologies, we analyzed the anterior prostate of the nonexposed F3 generation. The anterior prostates from the F1 and F3 generations were analyzed when the animals were 2 months old. For most of our experiments, we used a minimum of 4 biological replicates derived from independent, nonrelated treated mothers. For morphology evaluations, male progeny from a minimum of 8 nonrelated, treated females were used for each group; in total, 80 animals were analyzed.

### Anterior Prostate morphology analyses

To study anterior prostate morphology, anterior prostates from F1 and F3 control mice and the corresponding treated groups were fixed in Bouin’s solution for 24 h, washed in PBS and 70% ethanol, dehydrated, and embedded in paraffin. Five-micron sections of the entire prostate were cut, and the sections were placed on glass slides. The sections were deparaffinized and stained with H&E. Pictures were taken with a digital slide scanner (NanoZoomer, HAMAMATSU) and analyzed via NDPview software (v2.7.25) by two independent researchers. We analyzed 43 prostates in F1 and 37 in F3, for a total of 80 individual anterior prostates.

### Immunofluorescence of anterior prostate tissue in paraffin sections

For immunostaining, the paraffin blocks of the anterior prostates from the F1 and F3 control and treated groups were used. The sections were cut with a microtome (Microm HM 355 S) at a thickness of 5 μm. The sections were deparaffinized and rehydrated, and the epitopes were unmasked in 0.01 M citrate buffer, pH 6, at 80°C for 45 min. The slides were blocked and incubated with rabbit anti-PHH3 (1:500, 07--327, Millipore) or mouse PCNA (1:500, ab29, Abcam) antibodies. The sections were incubated with primary antibodies in PBS-T overnight at 4°C in a humidified chamber. After being washed in PBS-T, the sections were incubated with an appropriate fluorescent secondary antibody (1:100; AlexaFluor from Invitrogen) for 1 h in a humidified chamber at room temperature. The sections were counterstained with the Vectashield solution (Eurobio Scientific, France). The images were taken via an AxioImager microscope equipped with an AxioCam MRc5 camera and AxioVision software version 4.8.2 (Zeiss, Le Pecq, France) with a 5X or 40X objective lens using the same exposure time (DAPI, 350/442; Alexa Fluor 488, 488/525; Alexa Fluor 594, 594/617). The images were left unprocessed before analysis. For PCNA and PHH3, we quantified positive cells per area and analyzed a minimum of 6 images per slide from at least 6 different anterior prostates via ImageJ software. The data were averaged and are presented as the relative fluorescence compared with that of the control ± SD. The combined total corrected fluorescence (CTCF) was calculated as a subtraction from the integrated density of the mean intensity of the background area multiplied by the area. The mean intensity of the background was calculated for each image in the area without any cells. The data were averaged and are presented as positive cells per area compared with the control ± SD.

### Immunofluorescence of anterior prostate tissue in frozen sections

For immunostaining, anterior prostates from the F1 and F3 control and treated groups were embedded in optimal cutting temperature (OCT), and 7 mM sections were cut, dried, and stored at-80°C before use. The slides were fixed with 4% paraformaldehyde for 8 minutes at 4°C. The slides were quenched with 0.1 M glycine in PBS for 5 minutes and then washed 3 times with PBS. The slides were blocked with 4% BSA in PBS and incubated with rabbit anti-Ki-67 (1:500, ma5-14520, Invitrogen) and mouse BRD4 (1:500, sc-518021, Santa Cruz) or anti-mouse HDAC1 (1:500, sc-81598, Santa Cruz) and anti-rabbit KDM5A (1:500, ab194286, Abcam) antibodies diluted in DAKO (S2022, Agilent). The sections were incubated with primary antibodies overnight at 4°C in a humidified chamber. After 3 washes in 0.4% Kodak Photo Flo in PBS, the sections were incubated with an appropriate fluorescent secondary antibody (1:100; AlexaFluor from Invitrogen) for 1 h in a humidified chamber at room temperature. The sections were counterstained with the Vectashield solution (Eurobio Scientific, France). The images were taken via an AxioImager microscope equipped with an AxioCam MRc5 camera and AxioVision software version 4.8.2 (Zeiss, Le Pecq, France) with a 40X objective lens using the same exposure time (DAPI, 350/442; Alexa Fluor 488, 488/525; Alexa Fluor 594, 594/617). The images were left unprocessed before analysis. For Ki-67, we quantified positive cells per area and analyzed a minimum of 6 images per slide from at least 4 different anterior prostates via ImageJ software. For BRD4, KDM5A, and HDAC1, we used the lasso tool in ImageJ, drew the anterior prostate contour around epithelial cells, and measured the fluorescence. Combined total corrected fluorescence (CTCF) was calculated as a subtraction from the integrated density of the mean intensity of the background area multiplied by the area. The mean intensity of the background was calculated for each image, and we used the area without any cells. The data were averaged and are presented as the relative fluorescence compared with the control ± SD or the number of positive cells per area compared with the control ± SD.

### RNA extraction and RT‒qPCR

For the analysis of the F1 and F3 anterior prostates, total RNA was extracted from the anterior prostates that were snap-frozen following dissection via the RNeasy plus mini kit (Qiagen) according to the manufacturer’s instructions. The kit includes DNA elimination step, additionally RNA was treated with RNase-Free DNase Set (79254, Qiagen). For RT-qPCR analysis, 6 biological replicates from control and F1-and F3-generation males were used. Reverse transcription was performed with 1 µg of RNA via the iScript Reverse Transcription Kit (Invitrogen) according to the manufacturer**’**s instructions. The resulting cDNA was diluted 10 times and used for quantitative PCR. qPCR was performed using a Bio-Rad 384 plate machine with iTaq Universal SYBR Green Supermix (Bio-Rad, 1725124). Ct values for *Rpl37*, a housekeeping gene, were used for normalization via CFX manager software provided with the Bio-Rad 384 plate machine. The sequences of primers used for qPCR are shown in Table S1. The data were analyzed and are presented as the mean values of the fold change (FC) compared with the control. A nonparametric Mann‒Whitney test was applied to assess the statistical significance.

### Histone purification and western blot analysis

Protein samples from F1 and F3 mouse of the anterior prostates were prepared via the EpiSeeker Histone Extraction Kit (Abcam, 113476) according to the manufacturer’s protocol. Briefly, mouse prostates were homogenized via a TissueLyser (Qiagen) and centrifuged at 900 × g for 5 min. The pellets were resuspended in lysis buffer and left on ice for 30 min. After centrifugation, the supernatant fractions containing acid-soluble proteins were transferred to new tubes, and balance-DTT buffer was added. The protein concentrations were determined via the Pierce™ 660 nm Protein Assay (ThermoScientific, France). Five micrograms of protein were run on a 4-15% Mini-Protean precast polyacrylamide gradient gel (Bio-Rad, USA) for 1 h, and the proteins were transferred onto ImmobilonPSQ membranes (Millipore, France) via an electroblotter system (TE77X; Hoefer, USA) for 2 hours. Subsequently, blocking was carried out via the addition of 5% skim milk in PBS-Tween 20 (0.05%) for 1 hour. Proteins were detected via rabbit polyclonal anti-trimethyl-histone H3K4me3 (Millipore, 07-473; 1:10,000 dilution) or rabbit polyclonal anti-hyperacetylated histone 4 (Penta, Millipore, 06-946; 1:10,000 dilution). The signal intensities of histone modifications were normalized against the band intensities from the red Ponceau-stained membrane. The data are presented as normalized signals compared with the control +/-SD. The uncut original WB images are provided in supplementary Figure S1 and Figure S2.

### Chromatin immunoprecipitation and ChIP‒qPCR

We performed ChIP using rabbit polyclonal antibodies against H3K4me3 (07-473, Millipore). Equal amounts of material (∼anterior prostate from one mouse) were used and incubated in 1% paraformaldehyde solution for 10 minutes to crosslink proteins to DNA. One hundred microliters of 1.25 M glycine were added to each sample to quench the unbound paraformaldehyde. The samples were centrifuged, and 1 mL of PBS and two metal beads were added to the pellet, which was homogenized via TissueLyser (Qiagen). The samples were subsequently filtered in a cell strainer, and the resulting solution was pelleted and resuspended in the following buffer: 0.25% (vol/vol) Triton X-100, 10 mM EDTA, 0.5 mM EGTA, and 10 mM Tris (pH 8). The samples were centrifuged at 1100 × g for 5 minutes at 4°C, and the pellets containing the cells were resuspended in 300 μL of SDS lysis buffer (1% (wt/vol) SDS, 10 mM EDTA, and 50 mM TrisCl, pH 8) in the presence of a protease inhibitor and 10 mM DTT and incubated for one hour at RT. Chromatin was sonicated in SDS lysis buffer at 60% amplitude for 8 minutes (20 seconds on, 20 seconds off) via a Qsonica 700 sonicator (Q700--110, Newtown, Connecticut, USA) supplied with a 431C2 cup horn; these parameters allowed us to obtain ∼300 bp chromatin fragments. After sonication, the samples were centrifuged at 12800 rpm for 10 minutes at 4°C, and the supernatant containing the sonicated chromatin was transferred and diluted in 1.7 mL of the following buffer: 0.01% (1.1% (vol/vol) Triton X-100, 1.2 mM EDTA, 16.7 mM TrisHCl, and 167 mM NaCl. A solution containing 20 μL of Dynabeads (10002D, Invitrogen) and 0.7 µL of anti-H3K4me3 antibody (07-473, Millipore) was added to the sample tubes and incubated overnight at 4°C. Before the antibody and Dynabeads were added, 10 μL of each sample was collected as “Input samples” (starting material). After overnight incubation with Dynabeads and the antibody of interest, the beads were washed 5 minutes each, with the following four buffers: (1) Low salt buffer: 0.1% (wt/vol) SDS, 1% (vol/vol) Triton X-100, 2 mM EDTA, 20 mM TrisHCl, 150 mM NaCl; (2) High salt buffer: 0.1% (wt/vol) SDS, 1% (vol/vol) Triton X-100, 2 mM EDTA, 20 mM TrisCl pH 8, 500 mM NaCl; (3) LiCl buffer: 0.25 M LiCl, 1% (vol/vol) Igepal, 1 mM EDTA, 10 mM TrisCl, pH 8, 1% (wt/vol) deoxycholic acid; (4) TE buffer (two washes). Following the washing steps, the beads were resuspended two times in 50 µL of 1% (wt/vol) SDS and 0.1 M NaHCO3 (pH=9) and incubated at 65°C for 15 minutes to elute the precipitated chromatin from the beads. The eluted chromatin was subsequently reverse crosslinked by adding 9 µL of 5 N NaCl and incubating at 65°C for 4 hours. The proteins were subsequently removed by adding 1 µL of 20 mg/ml proteinase K and incubating the samples for 1 hour at 45°C. The precipitated DNA was purified via a MiniElute Reaction Clean-Up Kit (Qiagen), and the DNA concentration was measured via the QuantiFluor dsDNA system (Promega). A minimum of ∼10 ng of DNA was obtained.

Equal amounts of precipitated DNA and input samples were used for the qPCR analysis. Quantitative PCR was performed using 0.4 ng of immunoprecipitated or input DNA and 6 biological replicates. Normalized expression values were calculated with the CFX Manager program using a region located far from the promoter as a reference gene, and we used a region in *Gapdh* for H3K4me3-ChIP normalization. The primers used in this study are listed in Table S2. The enrichment of each target in the precipitated DNA was evaluated by calculating the ratio between the average of the normalized ChIP DNA copies and the average of the normalized DNA copies in the inputs.

## Statistical analyses

We used the minimum number of animals according to the requirements of the EU Ethics Committee. To assess the statistical significance of epithelial hyperplasia lesions in the prostates of the mice, we performed a Chi-square test. Given that the number of biological replicates in each experiment was relatively low, we performed a nonparametric Wilcoxon-Mann-Whitney test to assess the statistical significance of the results of the qPCR experiments, immunofluorescence, and western blot quantification.

### Results Experimental design

In this study, we analyzed the transgenerational effects of the neonicotinoid thiacloprid on the prostate. Pregnant outbred Swiss mice were exposed during the embryonic period from E6.5 to E15.5 for a total of 10 days. The thiacloprid dose used in this study was 6 mg/kg body weight/day. The chosen dose was approximately 4 times lower than the LOAEL established by the EPA for thiacloprid. The LOAEL dose for developmental neurotoxicity of thiacloprid is 25.6 mg/kg/day based on decreased preweaning and postweaning body weights in both sexes and delayed sexual maturation in males (EPA). A schematic of the experiments is shown in Figure S3. F1 and F3 progeny mice were studied at the age of 2 months; at this age, the prostate is functional, and the mice are sexually mature. Testosterone (T) in the blood serum was measured in a previous study; the level of T was not significantly modified in F1 males but was significantly decreased in F3 males ^23^. In this study, we analyzed anterior prostate morphology by staining paraffin sections with hematoxylin and eosin in forty-three F1 and thirty-seven F3 males; in total, 80 male anterior prostates were analyzed. We analyzed the proliferation marker PCNA and phosphorylated histone H3 at serine 10 (PHH3) in prostate paraffine sections. The chromatin histone modifiers HDAC1 and KDM5A and the oncology markers Ki-67 and BRD4 were analyzed via immunofluorescence in frozen sections. We analyzed the histone marks H3K4me3 and H4Ac by purifying the histones and analyzing them via western blotting. We determined the expression of the genes known to encode proteins important for prostate function and development via RT-qPCR. We analyzed histone H3K4me3 levels at the promoters of the genes via chromatin immunoprecipitation followed by qPCR. Finally, we analyzed the DNA methylation state of the genes important for prostate development via our previously published datasets.

### Direct exposure to Thiacloprid induces epithelial hyperplasia in the anterior prostate

To identify the effects of Thiacloprid on prostate morphology, paraffin-embedded prostate sections were stained with hematoxylin and eosin (H&E), and images were acquired using a digital NanoZoomer scanner. In our analysis, epithelial hyperplasia was identified in the anterior prostate, characterized by regions with a high density of rapidly proliferating epithelial cells forming so-called pseudostratified layers. The cells in the prostate exhibiting epithelial hyperplasia (EPH) display a stratified arrangement with dark, prominent nuclei and nucleoli. Representative images of control prostates and those exposed to epithelial hyperplasia in our study are shown in Figure 1a. In normal prostates, we observe irregularly arranged monolayer cells surrounding the lumen. In samples with epithelial hyperplasia prostate, regions of densely packed, cylindrical cells with dark nuclei and prominent nucleoli form pseudolayers. The percentage of epithelial hyperplasia prostates was analyzed in both F1 and F3 generations. In this study, the epithelial hyperplasia phenotype was observed in 5.9% of control group samples (1 out of 17 prostates) and in 27% of the F1-exposed group samples (7 out of 26 prostates) (p = 0.1, Chi-square test). In the F3 group, 7.1% of control prostates (1 out of 14) and 13% of F3-derived prostates (3 out of 23) exhibited the epithelial hyperplasia phenotype (p = 0.6, Chi-square test) (Figure 1b). Thus, our analysis revealed that gestational exposure to thiacloprid tended to increase the occurrence of hyperplasia in directly exposed F1 males, whereas no significant increase was detected in indirectly exposed F3 males.

**Figure 1.**
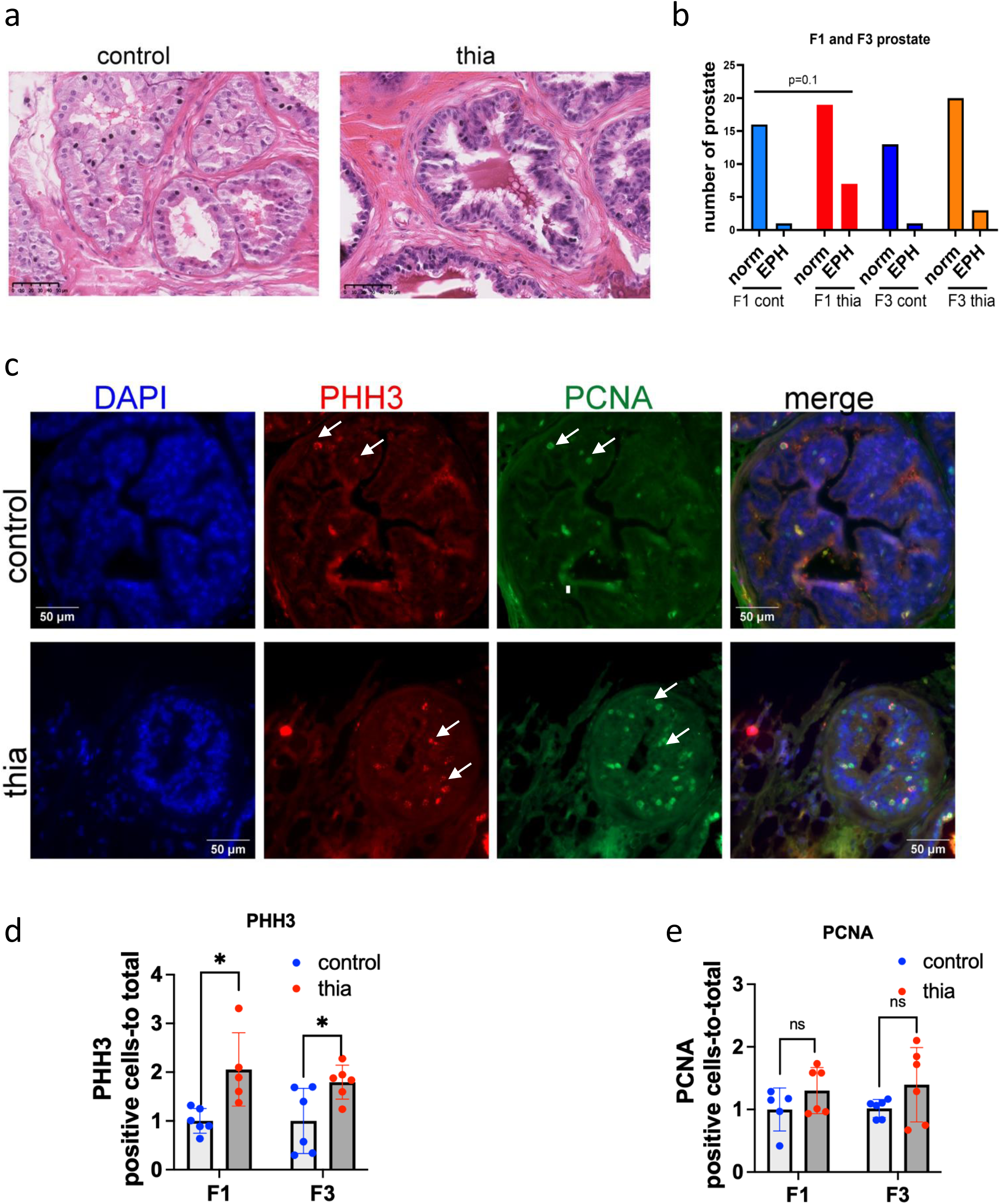
The prostate of these male mice has a greater number of epithelial hyperplasia and an increase in the expression of proliferation markers. (a) Representative images of prostates from control (left) - and *thia*-derived (right) mice. The prostate was embedded in paraffin, and 5 mM sections were stained with H&E. (b) Quantitative analysis of the prostate, F1: n= 17; control, n=26; F3, n=14; control, n=23. The exact p-value is indicated at the top of the graph, Chi-square test. (c) Representative images of the prostate coimmunostained by PCNA and phosphorylated histone H3 at Serine 10 (PHH3). (d) Quantitative analysis of PHH3-positive cells and (e) PCNA-positive cells. The PHH3 and PCNA plots in the figure represent the average values of the positive cells per area +/-standard deviation. F1, n=6 control; n=6 treatment; F3, n=6; control, n=6; *p<0.05; Mann‒Whitney test.

### Proliferation marker analysis revealed an increase in a marker of mitosis

To determine the capacity of prostate cells to proliferate, we carried out a marker analysis. We chose to analyze the expression of PCNA, a marker of cells that are in the early G1 phase and S phase of the cell cycle. PCNA helps in DNA repair during DNA replication ^24^. The level of histone H3 phosphorylation on serine 10 (PHH3) is correlated with chromosome condensation; thus, the phosphorylation state of Ser10 is a marker of mitosis ^25^. We prepared paraffin sections and coimmunostained them with antibodies against both of these markers as described in the Methods section. Our analysis revealed that both of these markers were localized in the nucleus; PCNA was detected throughout the nucleus, and PHH3 appeared as several bright dots in the nuclear periphery (Figure 1c). For both antibodies, we counted several positive cells per prostate area. The PHH3 marker significantly increased in both the F1 and F3 prostates by 2.1-and 1.8-times, respectively (Figure 1d). The quantitative analysis of PCNA revealed increases in F1 and F3 of 1.3-and 1.4-times, respectively (Figure 1e); however, the changes did not pass a significant cutoff threshold. Thus, our analysis of the mitosis markers revealed an increase in the prostates of the exposed mice, which is consistent with the tendency to increase the epithelial hyperplasia in F1 males.

### The oncogenesis marker analysis revealed an increase in F1 prostates

Since we observed an increase in the expression of mitosis markers, we verified whether cells could have alterations in oncogenesis markers. We analyzed the protein expression of the marker Ki-67 (also known as MKI67), which is a cellular marker for proliferation ^26^. Ki-67 remains active during the G1, S, G2, and M phases of the cell cycle ^27^ and is also an accepted hallmark of oncogenesis. We also include an analysis of bromodomain-containing protein 4 (BRD4), which is an epigenetic reader that regulates gene transcription through binding to acetylated chromatin via bromodomains ^28^. This marker is also abundant in cancer ^29^. To proceed with the experiment, we prepared frozen anterior prostate sections, as Ki-67 and BRD4 antibodies work better in frozen sections. We performed immunostaining in the F1 and F3 prostates as described in the Methods section. Ki-67 was highly intense in some cells, and BRD4 was detected throughout the nucleus (Figure 2a). We analyzed Ki-67-positive cells per prostate area. In F1, the number of Ki-67-positive cells increased significantly by 2.1-times, but no significant changes were detected in F3 (Figure 2b). In F1, BRD4 tended to increase 1.6-times (p=0.057), but no significant increase was detected in F3 (Figure 2c).

**Figure 2.**
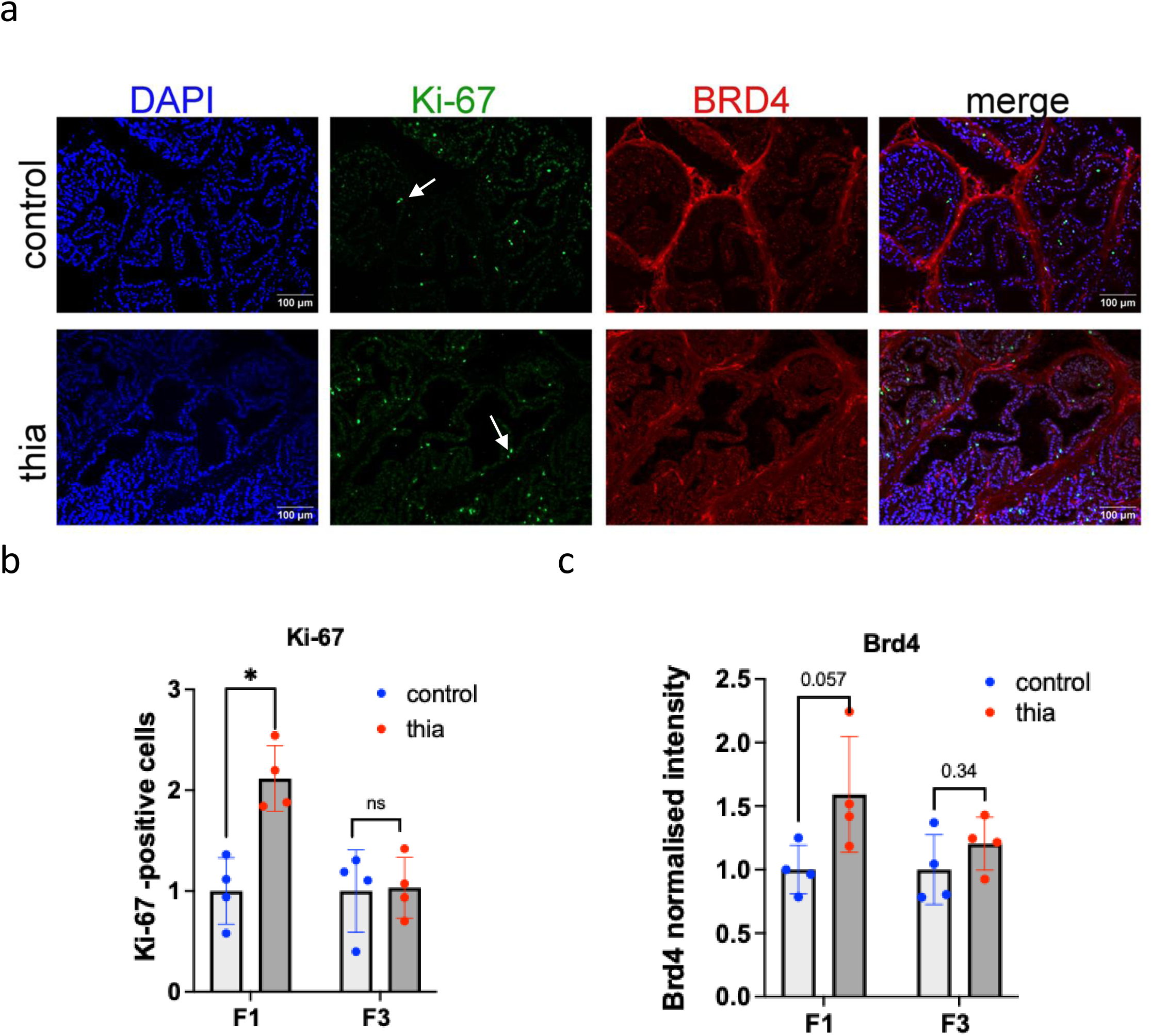
Oncology marker analysis of the prostates of male mice. (a) Representative images of prostates from control mice (top) and thia-derived F1 mice (bottom) immunostained with Ki-67 (green) and BRD4 (red). Quantitative analysis of the signal intensity of (b) Ki-67 and (c) BRD4. All plots in the figure represent average values +/-standard deviations. n=4 control, n=4 treatment in F1 and F3, *p<0.05, Mann‒Whitney test.

Our analysis revealed that gestational exposure to thiacloprid led to an increase in the proliferation marker Ki-67 in F1 but not in F3. Although BRD4 expression showed a trend toward an increase in F1, this change was not statistically significant. These results suggest that gestational exposure to thiacloprid may influence cell proliferation in F1, but these findings are not necessarily linked to oncogenesis, as Ki-67 is also associated with other biological processes. This increase in cell proliferation in F1 is consistent with a higher incidence of epithelial hyperplasia in these prostates, but it does not provide sufficient evidence for malignant transformation on its own.

### RNA expression analyses revealed increased expression of genes related to hormonal pathways, cellular proliferation, and epigenetic regulation

To gain insight into potential molecular mechanisms that are associated with the phenotypic changes that were observed in these derived males, we carried out RT-qPCR analysis using prostate tissue from F1 and F3 males. We performed a search for the genes that play a role in the prostate via the gene ontology program AMIGO, and we searched for a gene list that corresponds to the term “prostate development” (https://amigo.geneontology.org/amigo). Fifty-two genes were identified (Table S3). We added some targets on the basis of our previous research on prostate tissue ^30^. We extracted RNA from prostates and prepared cDNA as described in the Methods section. We analyzed the expression of the genes that encode genes related to transcription, hormonal signaling, and chromatin factors (Figure 3a-d). The housekeeping gene *Rpl37a* was used for normalization, and this gene was reproducible across all the samples.

**Figure 3.**
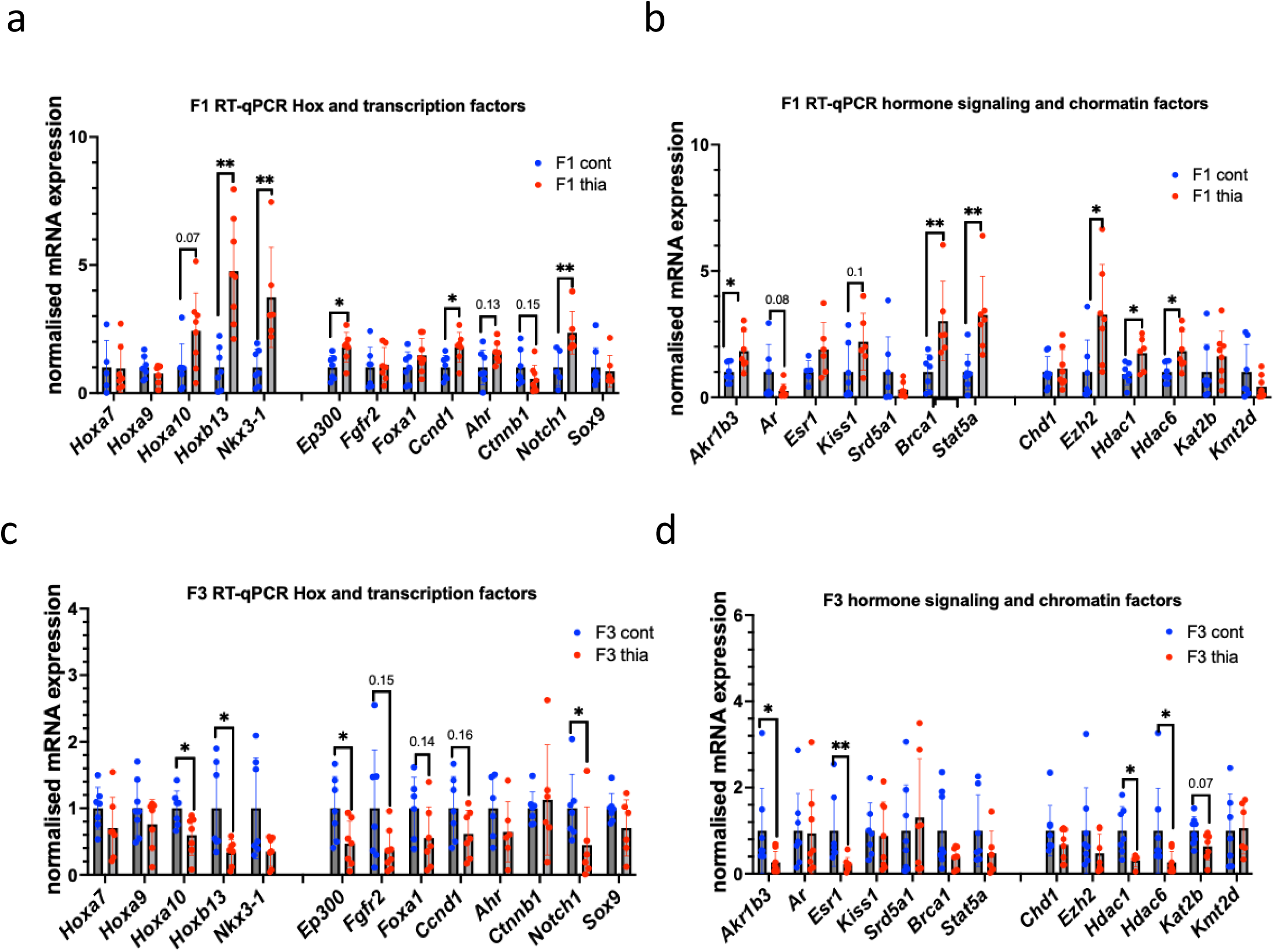
Effects of gestational thiacloprid exposure on gene expression in the prostate. (a-b) Gene expression analysis of the F1 prostate and (c-d) the F3 prostate. All plots in the figure represent average values of the fold change +/-standard deviation; n=6 for the control and treatment prostates from F1 and F3; *p<0.05, **p<0.01, Mann‒Whitney test.

In F1 anterior prostates, quantitative analysis revealed that *Hoxb13* and *Nkx3-1* expression increased by 4.8-and 3.7-fold, respectively. *Hoxa10* tended to increase expression by 2.4-fold. Both *Ep300* and *Ccnd1* increased expression by 1.8-fold, and *Notch1* increased expression by 2.4-fold. In addition, *Akr1b3*, *Brca1,* and *Stat5a* presented increases in expression of 1.8-, 3.0-and 3.3-fold, respectively. *Kiss1* tended to increase 2.2-fold. We also assessed the impacts of these factors on the expression of genes encoding chromatin factors. We determined that, in F1, the expression of *Ezh2, Hdac1*, and *Hdac6* increased by 3.2-, 1.7-and 1.8-fold, respectively. In contrast to the majority of the analyzed genes, the expression of the *Wnt* signaling pathway gene *Ctnnb1* tended to decrease 0.6-fold.

In F3, *Hoxa10, Hoxb13, Ep300,* and *Notch* expression decreased 0.6-, 0.3-, 0.5-, and 0.5-fold, respectively. *Fgfr2, Foxa1, and Ccnd1* tended to decrease 0.4-, 0.6-and 0.6-fold, respectively. In addition, *Akrb3, Esr1*, *Brca1, Hdac1* and *Hdac6* presented decreases in expression of 0.3-, 0.2-, 0.3-, 0.3-and 0.6-fold, respectively.

In summary, our analysis of RNA expression in the prostates of F1 and F3 offspring revealed that gestational exposure to thiacloprid altered the expression of genes associated with prostate morphology, cellular proliferation, chromatin factors, and genes encoding proteins of the hormonal signaling pathway. Most of the analyzed genes were upregulated in F1 and downregulated in F3.

### Analysis of the major regulatory histones revealed the global impact of thiacloprid on H3K4me3 levels in F1 and F3 prostates

Next, we analyzed the effects of thiacloprid on epigenetic regulation. To this end, we chose to investigate the effects of H3K4me3 on major regulatory marks. H3K4me3 marks are abundant in regions with open chromatin. We analyzed the H4 acetylation level because this marker is associated with chromatin compaction and it is abundant in cancer cells. We extracted histone proteins and performed western blot analysis. We revealed that H3K4me3 levels were increased in both the F1 and F3 prostates by 2.0-and 1.8-fold, respectively (Figure 4a-b). Histone H4 acetylation levels tended to increase 1.9-fold in F1 offspring (p=0.15). In contrast to H3K4me3, the H4 acetylation level tended to decrease 0.7-fold (p=0.11) in F3 (Figure 4c-d).

**Figure 4.**
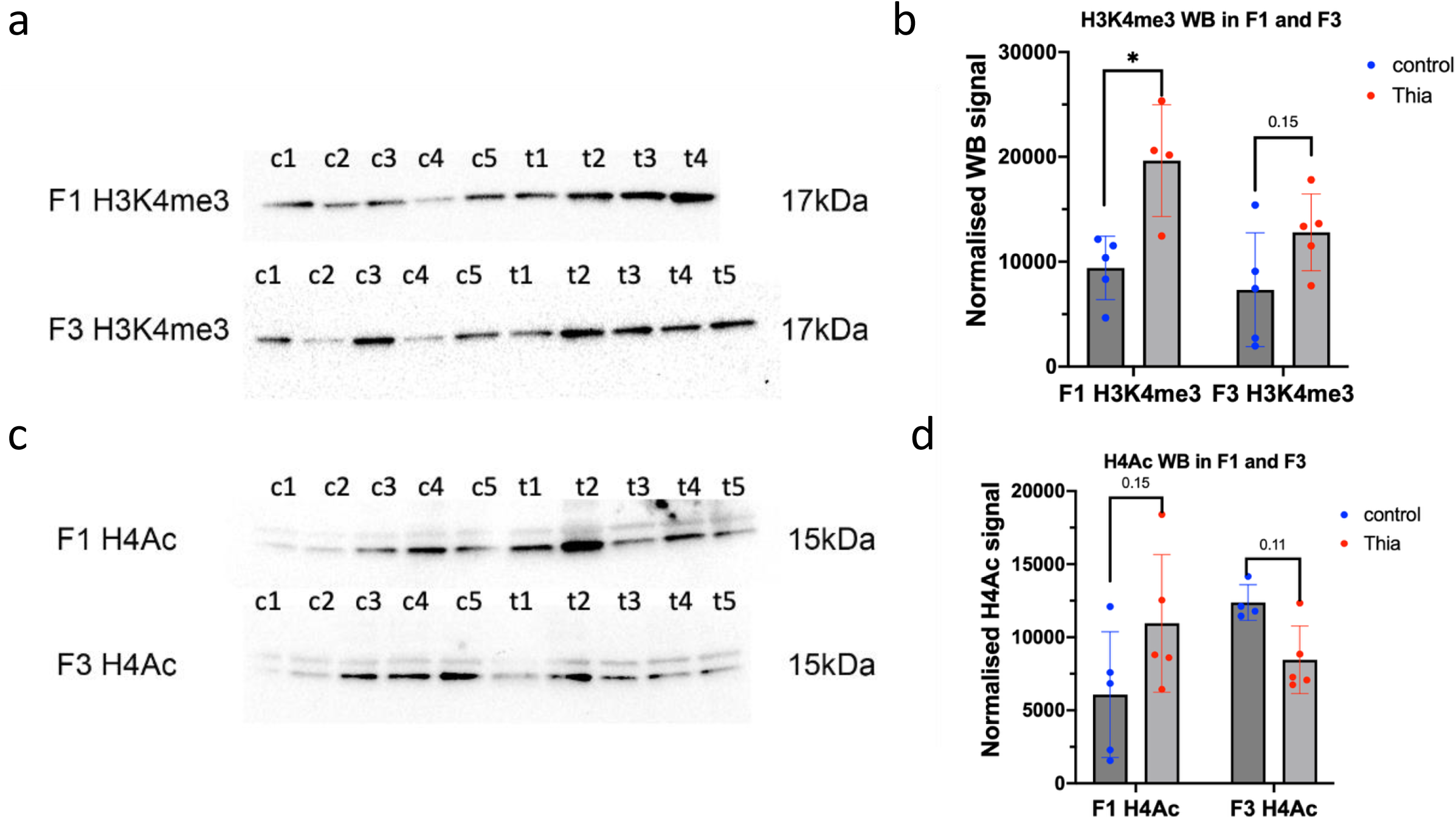
Analysis of purified histone proteins via WB. (a) Representative image of H3K4me3 c1--c4; c5 represents the control, and t1--t5 represent the treatment samples. (b) Quantitative analysis of H3K4me3 and (c) WB analysis of H4Ac by western blot. (d) Quantitative analysis of H4Ac; all plots in the figure represent average values +/-standard deviations. *p<0.05, nonparametric Mann‒Whitney test.

Thus, our analysis revealed that major regulatory marks changed, suggesting that exposure to thiacloprid has a global impact on the processes associated with these regulatory marks, on gene expression, and chromatin compaction.

### H3K4me3 analysis of prostate genes revealed increased occupancy in the prostate of F1 mice and decreased occupancy in F3 mice

Since H3K4me3 marker was altered globally in F1 and F3, we performed ChIP‒qPCR to confirm the role of this mark in gene expression changes. We designed primers at the regions of the genes that were chosen for RT‒qPCR analysis via the primer-blast program from ncbi.nih.gov and our previously published datasets of H3K4me3 in the prostate ^30^. As a control for the ChIP background, we used the region of the *Gapdh* gene, which is located distal to the promoter. We performed ChIP and analyzed the quantity of immunoprecipitated material compared with that of a starting material named Input, as described in the Methods section. The quantitative analysis revealed that the global histone H3K4me3 changes were consistent with the gene expression changes (Figure 5d). H3K4me3 occupancy at *Hoxa9* was elevated by 1.5-fold. H3K4me3 levels at the promoters of *Hoxa10* and *Hoxb13* tended to increase by 1.6-and 2.8-fold, respectively. *Ep300* and *Ccnd1* increased H3K4me3 levels by 1.8-and 1.9-fold, respectively. A significant increase in H3K4me3 was also detected at *Akr1b3* (2.2-fold), *Esr1* (3.3-fold), *Srd5a1* (2.0-fold), and *Stat5a* (2.4-fold). Chromatin factors have elevated levels of H3K4me3 at their promoters, specifically, at *Ezh2* (1.7-fold), *Hdac6* (1.4-fold), and *Hdac1* (3.4-fold). Notably, *Ctnnb1* decreased H3K4me3 by 0.5-fold, similar to the change in gene expression.

**Figure 5.**
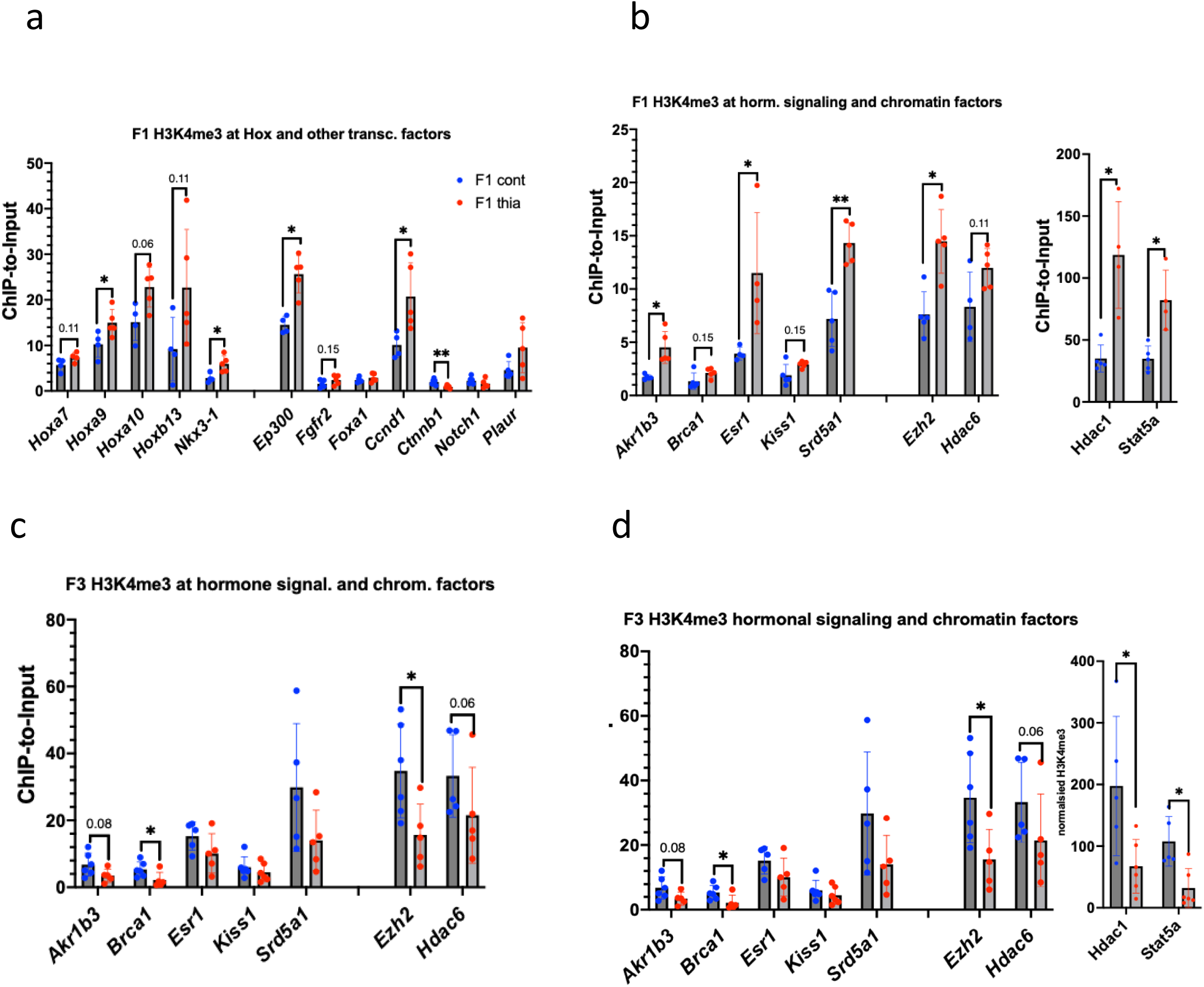
Effects of gestational thiacloprid exposure on H3K4me3 histone occupancy. (a-b) Analysis of H3K4me3 occupancy at the promoters of homeobox genes, genes encoding transcription factors involved in hormone signaling, and chromatin factors in F1 and (c-d) F3, F1 n=5 controls, n=6 treatment, and F3 n=5 controls, n=6 treatment. All plots in the figure represent average values +/-standard deviations. *p<0.05, Mann‒Whitney test.

In F3, similar to the gene expression changes, we detected a decrease in H3K4me3 at the promoters of the analyzed genes. Specifically, we noted significant decreases in *Hoxa9* (0.5-fold), *Hoxa10* (0.5-fold), *Hoxb13* (0.6-fold), and *Nkx3-1* (0.4-fold) expression. We determined that Ep300 tended to decrease (0.6-fold). The H3K4me3 levels at *Brca1* and *Ezh2* decreased 0.4-fold. *Akrb13, Esr1, and Hdac6* tend to decrease H3K4me3 by 0.5-, 0.7-, and 0.6-fold, respectively. *Hdac1* and *Sta5a* both decreased H3K4me3 levels by 0.3-fold.

In summary, our analysis of H3K4me3 showed that there are similar alterations in *thia*-derived males regarding gene expressions, suggesting that thiacloprid exposure induces epigenetic alterations that probably impact transcription factor binding. However, in F3, despite the increase in H3K4me3 histone protein levels, the vast majority of the analyzed genes presented a decreased level of H3K4me3, suggesting that some other regions, not analyzed by ChIP‒qPCR, could contribute to the global level of H3K4me3.

### Increased HDAC1 levels were detected in the F1 anterior prostate

Since class I HDAC levels are increased in anterior prostate cancers and their aberrant expression is correlated with decreased tumor suppressor activity and drug resistance ^31,32^, we analyzed HDAC1 levels in the F1 and F3 anterior prostates via immunofluorescence. Histone deacetylase 1 (HDAC1) is a KDM5A-dependent factor, and these two factors coregulate their downstream genes ^33^. KDM5A is also involved in the transcriptional regulation of *Hox* genes and plays a role in tumor progression ^34^; thus, we analyzed the protein level of KDM5A as well. We prepared frozen sections and fixed them with paraformaldehyde, and we performed immunofluorescence analysis of the intensity of these markers per area of prostate epithelial cells. Our analysis revealed that both of these markers have strong staining in the nucleus and are colocalized (Figure 6a). In F1, we detected a significant increase in the HDAC1 level of 1.4-fold (Figure 6b). In contrast to F1, we observed a slight but not significant decrease of 0.9-fold in HDAC1 in the F3 anterior prostate. KDM5A analysis revealed no significant changes in the prostates (Figure 6c).

**Figure 6.**
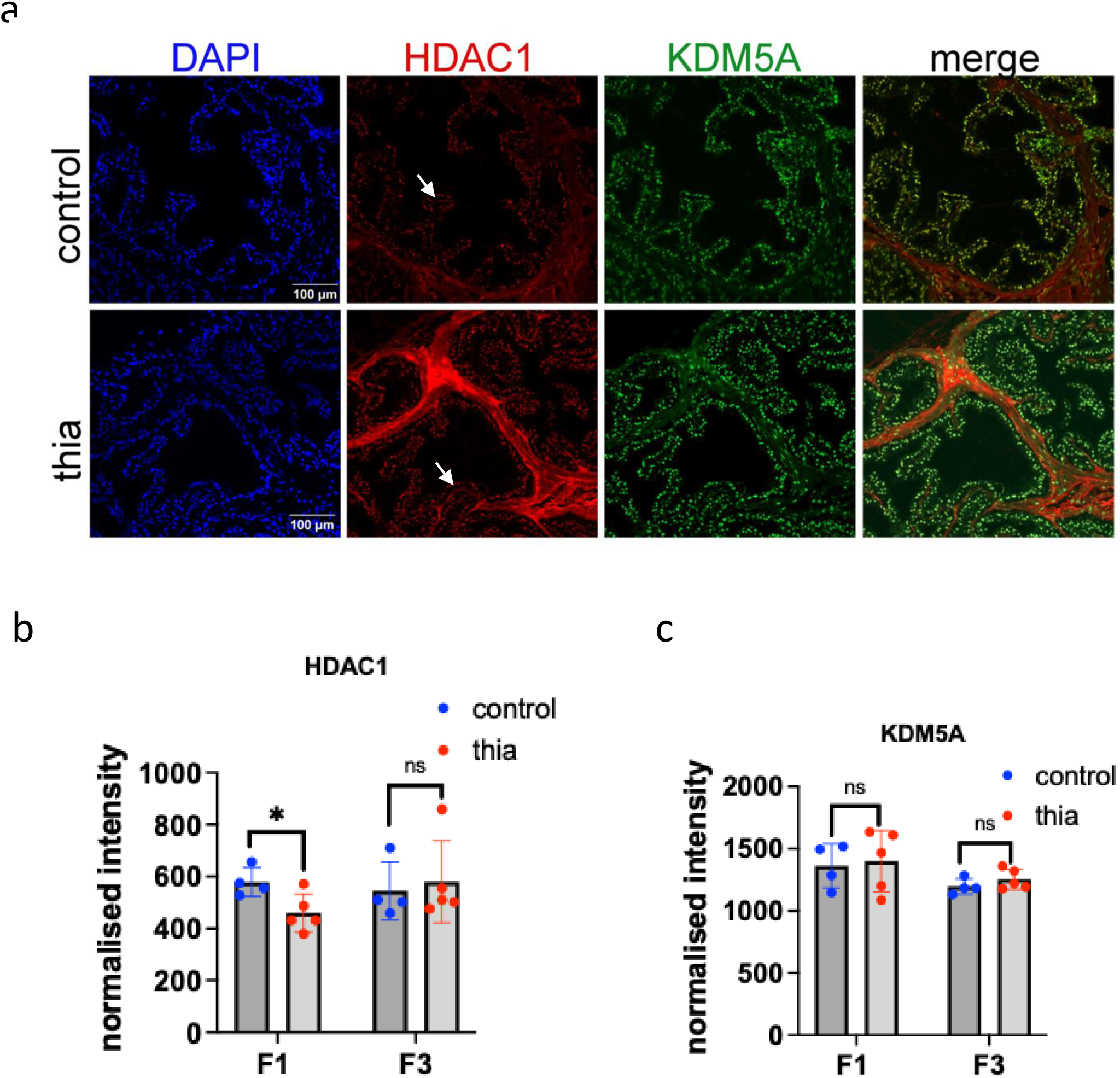
Chromatin marker analysis of the prostate of male mice. The mice were sacrificed at 2 months of age. (A) Representative images of prostates immunostained for KDM5A (green) and HDAC1 (red) from control (top) and treatment (bottom) mice. (B) Quantitative analysis of the signal intensity of the markers. All plots in the figure represent average values +/-standard deviations. n=4 control and treatment in F1 and F3, *p<0.05, Mann‒Whitney test. All plots in the figure represent average values +/-standard deviations.

In summary, we observed an increase in HDAC1 levels in the F1 anterior prostate, and this increase in HDAC1 intensity is consistent with elevated *Hdac1* gene expression and increased H3K4me3 levels at the promoter of HDAC1. The intensities of KDM5A in F1 and F3 as well as HDAC1 in F3 were not significantly modified.

### Sperm DNA methylation analysis revealed a potential impact of Hox regions on the toxicity of thiacloprid in prostate tissues

To reveal how the induced effects in the prostate of F1 are inherited by F3 males, we analyzed the sperm of F1and F3 male at 2 months old thiacloprid-derived males compared with those of the control. To this end, we reanalyzed our recent sperm DNA methylation data from ancestral thiacloprid exposure ^23^. We examined DNA methylation at *Hdac1* and determined that in the vicinity of ∼20 kb, there is a DNA methylation mark that tends to decrease in F1 and increase in F3 (Figure S4a-b), suggesting a possible impact of paternal sperm DNA methylation and histone H3K4me3 in the prostate at the *Hdac1* gene.

We also performed DNA methylation analysis of genes that are important for prostate development. We used the gene list determined from the AMIGO database as “prostate development” genes (Table S3). We verified the presence of DNA methylation marks near these genes in our sperm datasets from F1, F2, and F3 males and compared the samples from *thia-* derived males and the control. The analysis revealed that in sperm, several genes have alterations in DNA methylation in F1 and F3 males. For example, we observed DNA methylation marks in the region overlapping the *Hoxa3* gene (Fig. 7a). This region also overlaps with the superenhancer element mSE_04719, which is active in the embryonic brain. Notably, the region was found to have decreased DNA methylation in F1 and F2 sperm but increased methylation in F3 sperm. In addition to *Hoxa3*, some other regions were found to have altered DNA methylation in the sperm, including regions near the *Stk11, Ahr, Nkx3-1, Plaur, Tnc, Wdr77, Fgfr2, Eaf2, Igf1, Fem1b, Rln1, Ctnnb1,* and *Notch1* genes (Figure 7b). Notably, the changes in DNA methylation are in opposite directions in terms of gene expression changes and histone H3K4me3 marks (*Nkx3-1, Ahr,* and *Fgfr2*).

**Figure 7.**
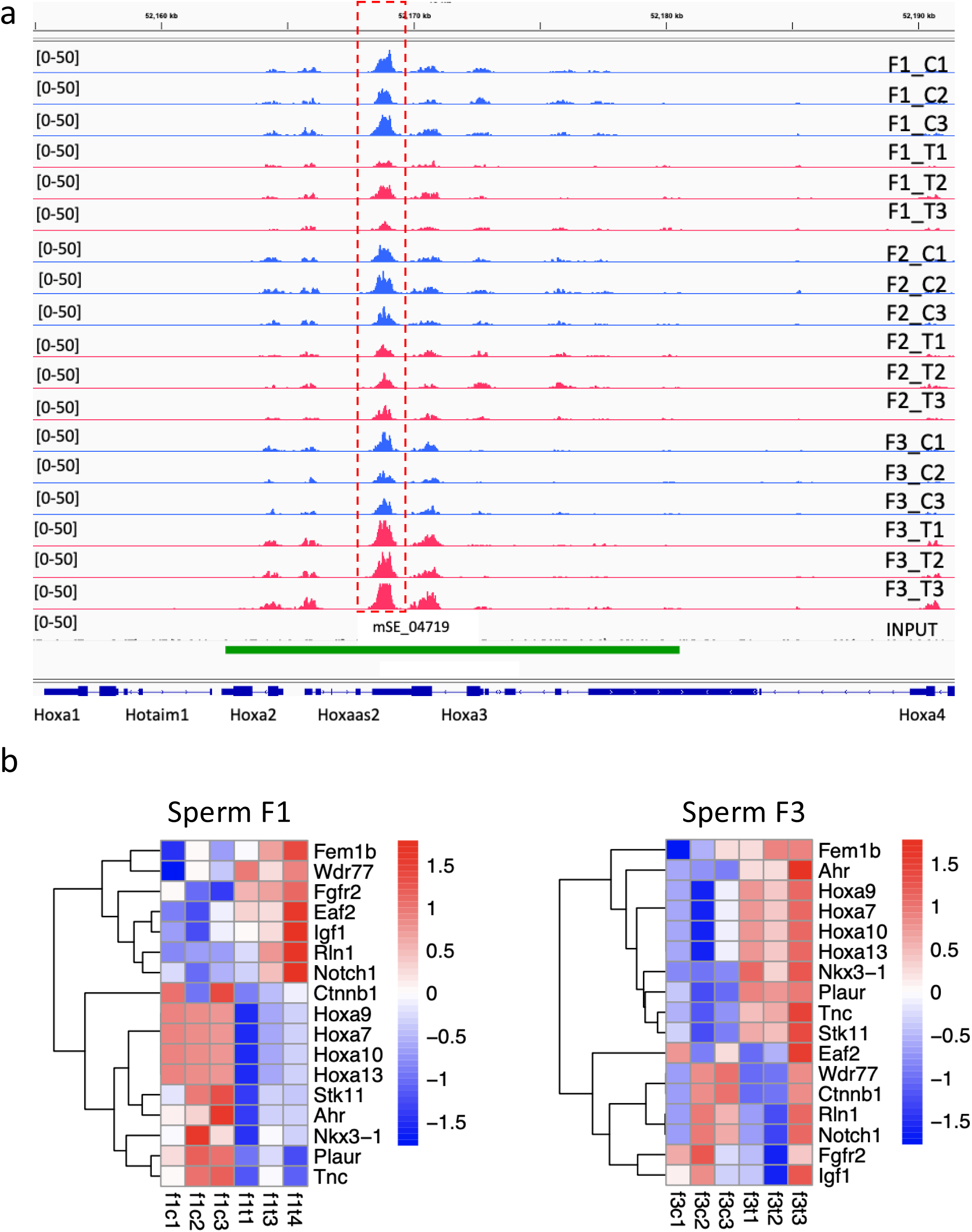
DNA methylation analysis of the genes that play a role in prostate development in the sperm of F1 to F3 mice. (a) Plots of differentially methylated regions overlapping *Hoxa3* gene; the signal range is indicated in brackets; the differential peak is marked by the red dashed box. (b) Heatmap of DMRs in the sperm of F1 and F3 males. MeDIP counts with FC>1.5 and FDR<0.1 were log-transformed and plotted in R via Heatmap; f1c1--f1c3 are F1 controls; f1t1--f1t3 are F1th-derived samples; f3c1--fc3 are F3 controls; and f3t1--f3t3 are F3th-derived samples. The sequencing analysis was performed using sperm DNA and a minimum of 3 replicates for each group from F1 to F3.

Our observations suggest that paternal sperm DNA methylation affects prostate histone marker alterations in exposed mouse progeny.

## Discussion

### Effects of thiacloprid on prostate morphology

We determined the tendency toward an increased epithelial hyperplasia number of prostates with epithelial hyperplasia and increased expression of mitosis (PHH3) in both F1 and F3 generation males. Our data are consistent with previous observations of other authors, who noted that endocrine-disrupting compounds such as BPA ^35^ or phthalates ^36^ could induce the appearance of epithelial hyperplasia in anterior prostate. In these studies, the exposure periods were gestational or early neonatal, suggesting that early development is critical for the establishment of cell lineages specific to the prostate. Indeed, alterations in developmental genes of the prostate could impact the normal development of the prostate in adult animals. The morphology and proliferation markers were less detectable in F3. Thus, most of the changes induced in the prostate in the directly exposed F1 generation were not inherited by F3. It is conceivable that under toxic exposure, some pathologies are induced. When the environment is no longer toxic, some epigenetic alterations are reversed as we observed in F3 males.

### Thiacloprid-induced epigenetic effects

In this study, we analyzed H3K4me3 by WB and ChIP‒qPCR. The H3K4me3 global level increased in both generations, which could be due to an increased level of mitosis resulting from a significantly increased level of PHH3, a marker of mitosis, in F1 and F3. A high level of H3K4me3 in the prostate was previously shown in our previous transgenerational study ^37^, and we observed a higher level of proliferation markers and an increased rate of epithelial hyperplasia. The increase in proliferation changes the chromatin structure to a more accessible configuration ^38^. H3K4me3 marks are recognized by many transcription factors and other chromatin readers ^39^; therefore, H3K4me3 marks create a platform for an increased transcription. For example, in the developing prostate, endocrine-disrupting compounds induce MLL1/KMT2A activation, which causes increased H3K4me3 levels in genes associated with prostate cancer, and importantly, the elevated expression of these genes persists in adulthood ^40^. Elevated H3K4me3 levels were detected in hormone-resistant prostate cancer HRPC^41^.

In contrast to H3K4me3, H4Ac tended to increase in F1 and decrease in F3 in our study, suggesting that exposure to *H4Ac* creates more accessible chromatin in F1 than in F3. Notably, although H4Ac tends to increase in F1 offspring, we observed that the HDAC1 protein was abundantly present in our F1 offspring. An abundance of HDAC1 was previously observed in malignant epithelial nuclei in prostate tissue ^42^. The differential HDAC expression in epithelial and stromal cells may play important roles in the progression of prostate cancer ^43^. We did not measure the HDAC1 level in stromal tissues, but it is possible that both stromal and epithelial cells could have differential HDAC levels.

We determined that DNA methylation of several genes essential for prostate development was altered. Indeed, *thia* exposure overlaps with the embryonic period of the somatic-to-germline transition when DNA methylation is erased and re-established de novo. The perturbation of the activity of DNA methyltransferases is induced by numerous environmental factors ^44;^ thus, it is conceivable that some alterations in DNA methylation in our study were established due to perturbation of *de novo* methyltransferases such as DNMT3A. Some of these alterations could be preserved by subsequent generations. However, most DNA methylation is erased after fertilization. Only limited regions resist reprogramming. For example, some IAP-containing regions avoid reprogramming ^45,46^. In our previous *thia-*exposed testis study, we showed that near the *Isl1* region, DNA methylation decreases in F1 and F3, and this region overlaps with the IAP retroelement. Recently, the role of ISL1 was demonstrated in prostate cancer. *Isl1* knockdown reduces androgen receptor (AR) activity and leads to reduced cell growth in castration-resistant prostate cancer ^47^, suggesting a potential role for *Isl1* in prostate proliferation in our study. The role of *Isl1* on pathology induced by thiacloprid was not evaluated in this study.

We also noted that alterations in our studies in F1 and F3 epigenetic marks had opposite effects. We cannot simply explain this phenomenon; we suggest that some unknown mechanisms compensate for the previously induced epigenetic alterations in F3 males. Similar opposite effects results were observed in our previous study in prostates exposed to chlordecone ^30^. We suggest that compensatory effects could be promoted during fertilization. The chromatin rearranged after fertilization to form a functional embryonic genome. In the paternal genome, chromatin nucleosome domains are re-established. The sequencing of early embryos revealed that the maternal genome started to reduce the size of the broad H3K4me3 peaks after fertilization, while paternal H3K4me3 marks are detectable at very low levels only in some regions ^48,49^, suggesting that H3K4me3 marks of both parental genomes rearranged in new embryos. A recent study revealed that in zebrafish, there are regions named“placeholders”, which contain the histone H2A variant H2A.Z and H3K4me1; these regions occupy all regions lacking DNA methylation in both sperm and embryos and reside at promoters encoding housekeeping and early embryonic transcription factors. Thus, other histone modifications, such as H2A.Z, may play a role in the rearrangement of histone and DNA methylation marks after fertilization and the establishment of new patterns in new embryonic tissues.

## Conclusion

Our study revealed that exposure to thiacloprid induces proliferation and is associated with epigenetic alterations in the sperm at genes important for prostate development.

## Limitations of this study

We analyzed murine prostates in young 2-month-old animals. We have to reduce the number of animals used in our studies to follow the requirements of the EU Ethics Committee; thus, the prostates in this study were recovered from other “testis” studies. Prostate pathologies normally appear at a later age; thus, we suggest that some alterations could be observable at a later age, but we did not study them.

## Ethics approval and consent to participate

All experimental procedures involving animals were authorized by the Ministry of National Education and Research of France (Number APAFIS#17473-2018110914399411 v3). The animal facility used for the present study is licensed by the French Ministry of Agriculture (agreement D35--238--19). All experimental procedures followed the ethical principles outlined in the Ministry of Research Guide for Care and Use of Laboratory Animals and were approved by the local Animal Experimentation Ethics Committee (C2EA-07). All methods were performed under the ARRIVE guidelines. The animals were euthanized by placing them in a carbon dioxide (CO2) chamber. All procedures were authorized by the Ministry of National Education and Research of France; the authorization number is #17473-2018110914399411 v3. All euthanasia procedures were performed according to annex IV of low 2013--118 issued by the Ministry of Agriculture, Food and Forestry of France on February 1, 2013.

## Consent for publication

Not applicable

## Availability of data and material

All data generated or analyzed during this study are included in this published article.

## Authors’ contributions

FS designed the research. OD, CDL, TMB, TD, CK and PYK performed the experiments. FS wrote the manuscript. All authors approved the final version of the manuscript.

## Supporting information

The uncut original WB images are provided in supplementary Figure S1 and Figure S2.

## Abbreviations

name: description
AHR: aryl-hydrocarbon receptor
AKR1B3: aldo-keto reductase family 1 member B
AR: androgen receptor
BRCA1: breast cancer 1, early onset
BRD4: bromodomain containing 4
CCND1: cyclin D1
ChIP: chromatin immunoprecipitation
CTCF: combined total corrected fluorescence
CTNNB1: catenin (cadherin associated protein), beta 1
DNMT3A: DNA methyltransferase 3A
EAF2 ELL: associated factor 2
EP300 E1A: binding protein p300
EPA: Environmental Protection Agency
EPH: epithelial hyperplasia
ESR1: estrogen receptor 1 (alpha)
EZH2: enhancer of zeste 2 polycomb repressive complex 2 subunit
FEM1B: fem 1 homolog b
FGFR2: fibroblast growth factor receptor 2
FOXA1: forkhead box A1
GAPDH: glyceraldehyde-3-phosphate dehydrogenase
GRRs: germline reprogramming responsive genes
H&E: hematoxylin and eosin
HDAC1: histone deacetylase 1
HDAC6: histone deacetylase 6
HOXA10: homeobox A10
HOXA3: homeobox A3
HOXA9: homeobox A9
HOXB13: homeobox B13
IAP: Intracisternal A particle
IGF1: insulin-like growth factor 1
IMI: Imidacloprid
ISL1: ISL1 transcription factor, LIM/homeodomain
KDM5A: lysine (K)-specific demethylase 5A
KI-67: marker of proliferation Kiel 67
KMT2A: lysine (K)-specific methyltransferase 2A
LOAEL: Lowest-observed-adverse-effect level
NKX3-1: NK3 homeobox 1
NOTCH1: notch 1
PCNA: proliferating cell nuclear antigen
PHH3: Phosphorylated Histone H3
PLAUR: plasminogen activator, urokinase receptor
RLN1: relaxin 1
RPL37: ribosomal protein L37
SGT: somatic-to-germline transition
STAT5A: signal transducer and activator of transcription
5A STK11: serine/threonine kinase 11
TNC: tenascin C

## Acknowledgements

The authors are thankful to the H2P2 platform of Rennes for assisting with tissue preparation for histology.

---

**Figure S1.** Schematic representation of the experiments. Pregnant outbred Swiss mice were treated from E6.5 to E15.5, with doses of 6 mg/kg/day or vehicle, and control mice received only vehicle (oil). F1 and F3 mice were sacrificed at the age of 2 months. The schema of breeding is described in the “Mouse treatment and dissection” section of the Methods section.

**Figure S2.** Uncut WB images

**Figure S3.** Ponceau red-stained membrane images

**Figure S4.** Fig S3. DNA methylation analysis of the *Hdac1* gene in the sperm of F1, F2, and F3 mice. (a) Plots of sequencing reads near the Hdac1 gene; the signal range is indicated in brackets. Normalized counts were averaged and plotted, f1c1-f1c3 are F1 controls, f1t1-f1t3 are F1 treatment samples, f2c1-f2c3 are F2 controls, f2t1-f2t3 are F1 treatment samples, 3c1-fc3 are F3 controls, and f3t1-f3t3 are F3 treatment samples. The sequencing analysis was performed using sperm DNA, with a minimum of 3 replicates for each group. The differential peak is marked by the red dashed box. (b) Quantitative analysis of Hdac1 counts in F1 (top) and F3 (bottom) graphs. The exact p-value is indicated at the top of the graph; Mann‒Whitney test.

**Table S1.** Oligonucleotides used for RT-qPCR

**Table S2.** Oligonucleotides used for ChIP-qPCR

**Table S3.** Biological process “prostate gland development”

