## Supplementary material for "Transgenerational effects induced by thiacloprid in Anterior prostate tissue are associated with alterations in DNA methylation at developmental genes": The uncut original WB images are provided in supplementary Figure S1 and Figure S2.

Univ. Rennes, EHESP, Inserm, Irset (Institut de recherche en santé, environnement et travail) - UMR_S 1085, F-35000, Rennes, France

**Supplementary information**

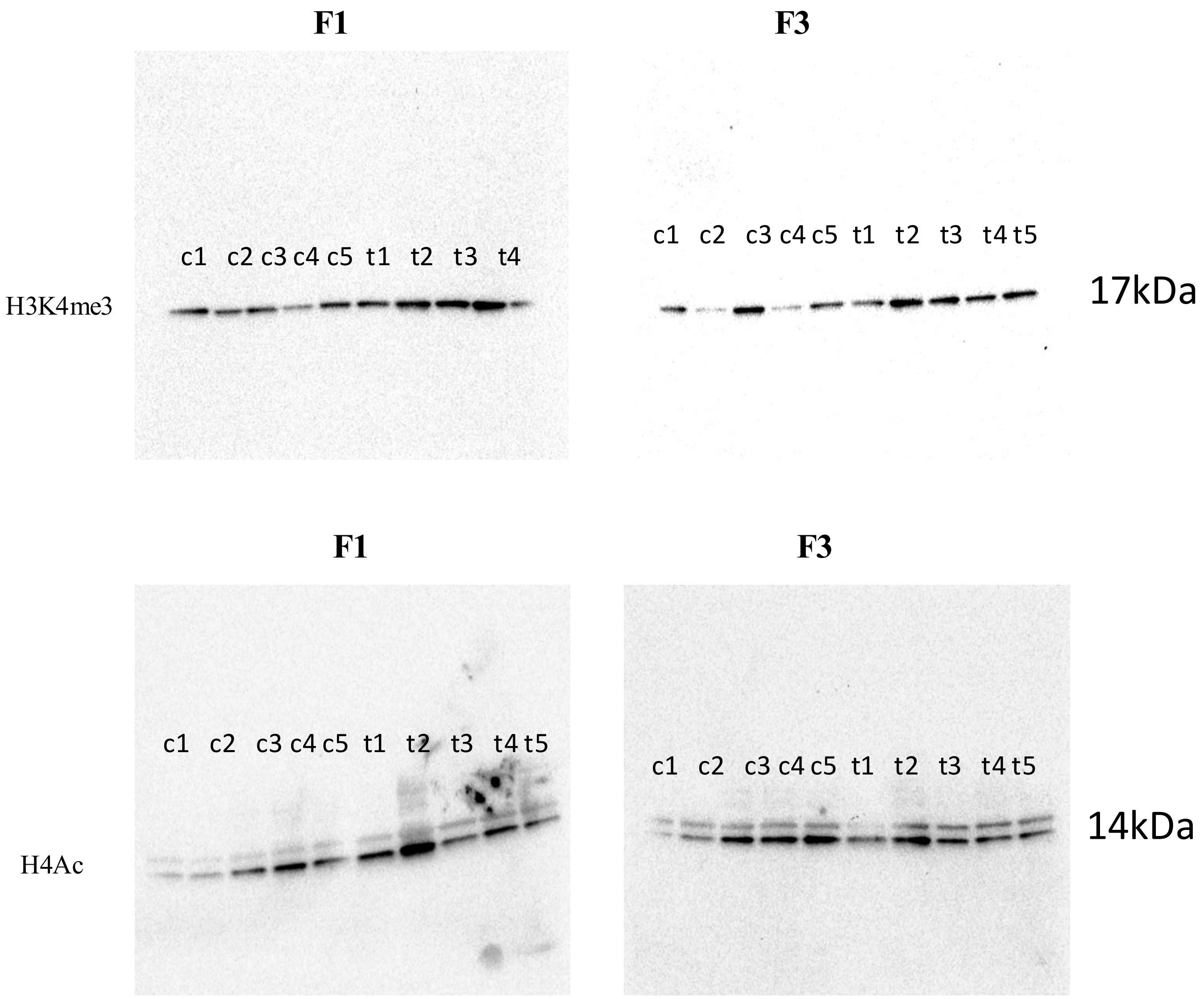


**Figure S1.** Uncut WB images

**F1** **F3**

H3K4me3

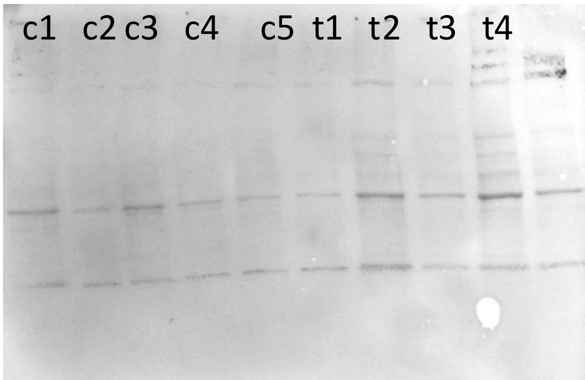


hist

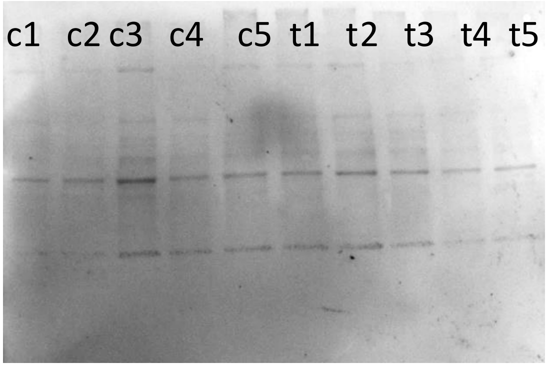


hist

**F1** **F3**

H4Ac

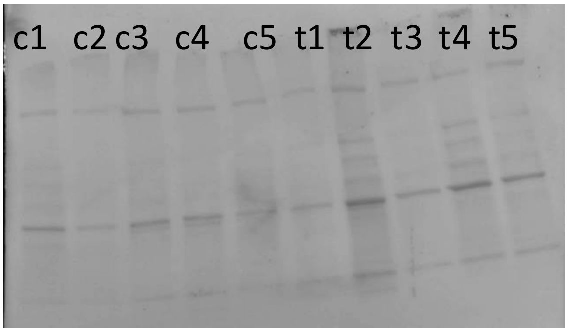


hist

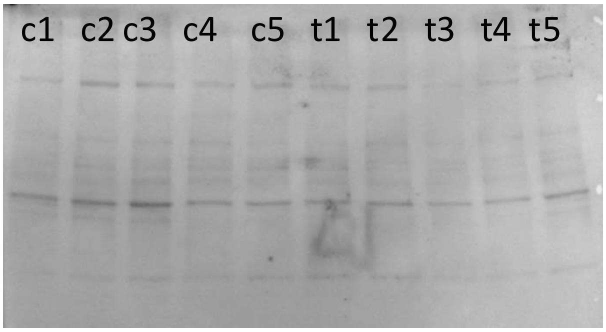


hist

**Figure S2.** Ponceau red -stained membrane images

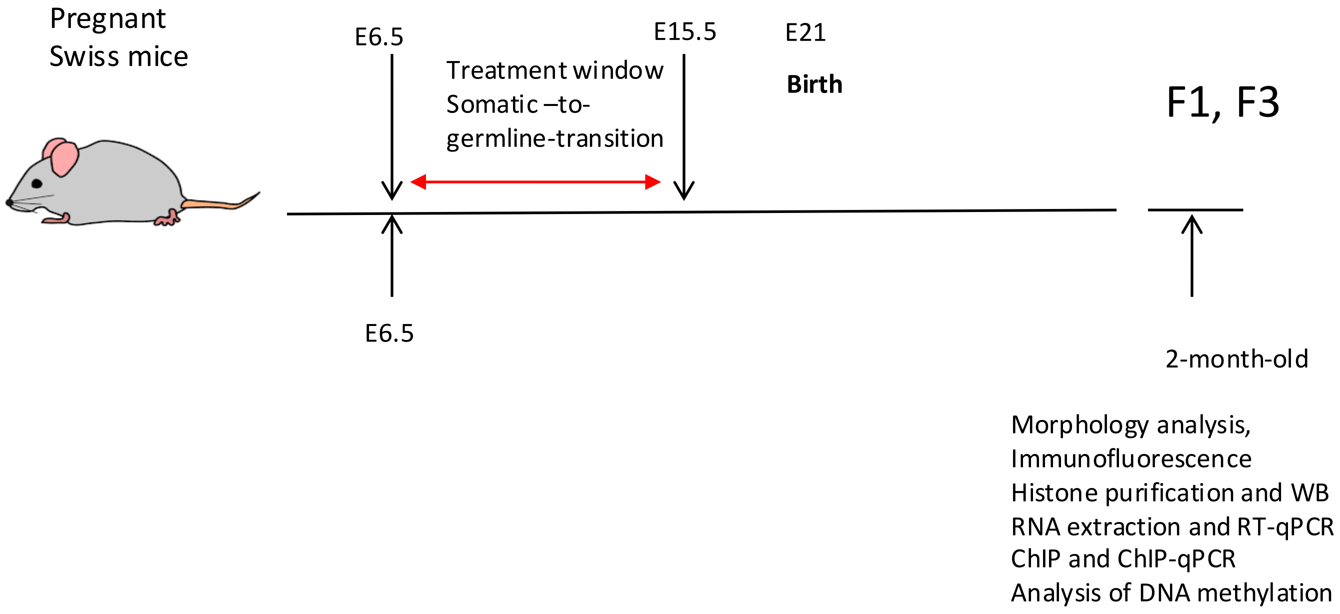


**Figure S3.** Schematic representation of the experiments. Pregnant outbred Swiss mice were treated from E6.5 to E15.5, with doses of 6 mg/kg/day or vehicle, and control mice received only vehicle (oil). F1 and F3 mice were sacrificed at the age of 2 months. The schema of breeding is described in the “Mice treatment and dissection” section of the Methods section.

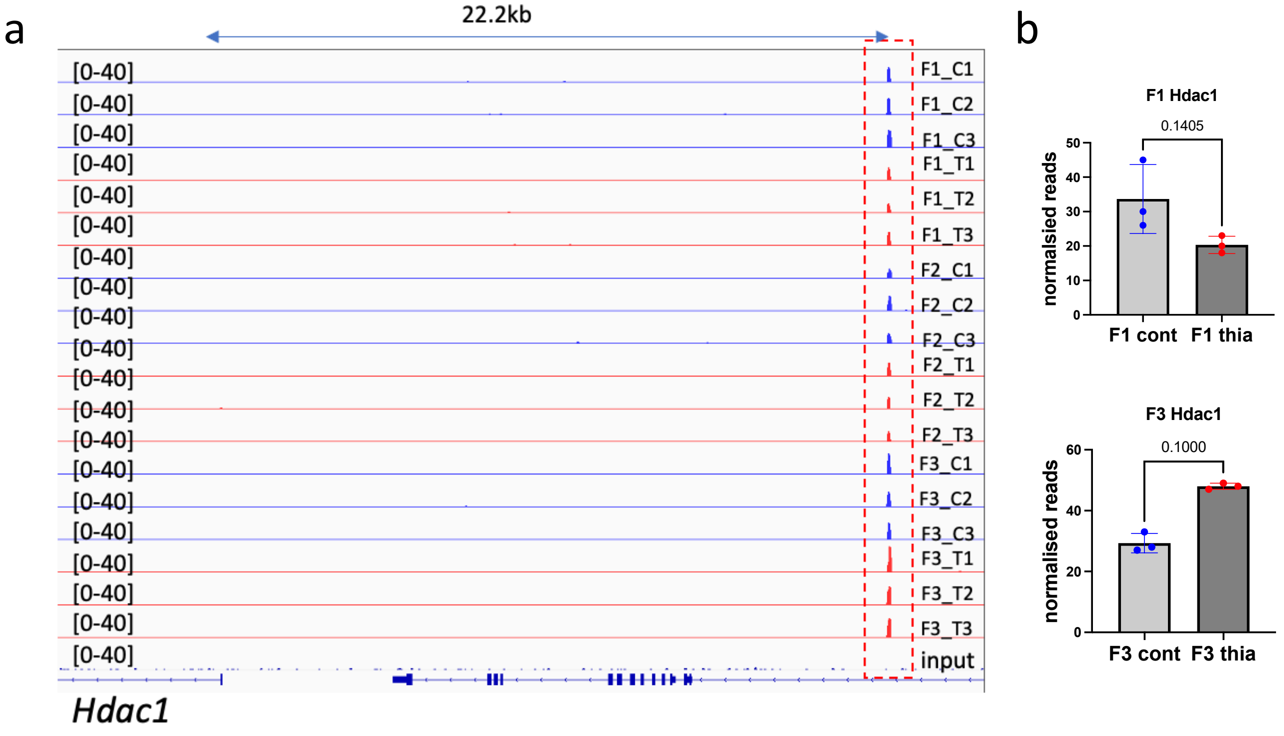


**Figure S4.** DNA methylation analysis of the *Hdac1* gene in the sperm of F1, F2, and F3 mice. (a) Plots of sequencing reads near the Hdac1 gene; the signal range is indicated in brackets. Normalized counts were averaged and plotted, f1c1-f1c3 are F1 controls, f1t1-f1t3 are F1 treatment samples, f2c1-f2c3 are F2 controls, f2t1-f2t3 are F1 treatment samples, 3c1-fc3 are F3 controls, and f3t1-f3t3 are F3 treatment samples. The sequencing analysis was performed using sperm DNA, with a minimum of 3 replicates for each group. The differential peak is marked by the red dashed box. (b) Quantitative analysis of *Hdac1* counts in F1 (top) and F3 (bottom) graphs. The exact p-value is indicated at the top of the graph; Mann–Whitney test.

### Table S1. Oligonucleotides used for RT-qPCR

| gene | forward | reverse | biological process |
|---|---|---|---|
| Ahr | AATCCCACATCCGCATGATTAAGAC | TGAGTGGCGATGATGTAATCTGGT | prostate gland development |
| Akr1b3 | CAAACCTTCATCCACTAGTTGTTCC | GGCCCGACTATTTCCCACTG | enables aldose reductase (NADPH) activity |
| Ar | GTCCTTCACTAATGTCAACTCCA | CCACTGGAATAATGCTGAAGAG | activation of prostate induction by androgen receptor signaling |
| Brca1 | CCCAAAGAAGTAATGACCGTG | GCTAACTATCCACTTTCCTCCTG | regulation of DNA damage checkpoint |
| Ccnd1 | GGACGTCGTCAGGAGCAC | ACCGACGTGCGAGATGTG | G1/S transition of mitotic cell cycle |
| Chd1 | TCATAAACCAACACAGTAATTGCC | GTTGGGATAATAGACCTTGCGT | chromatin organization |
| Ctnnb1 | TCAGTGCAGGAGGCCGAGG | TCCAACTCCATCAGGTCAGCTTG | Wnt signalling |
| Ep300 | CCTTCCACTCCGCTTTCTCA | ACCTTTAGCCTCCTTTGTATCCTC | animal organ morphogenesis |
| Esr1 | CACGTTTCTGTCCAGCACCTTGAAGT | AGAGATGCTCCATGCCTTTGTTACTCA | estrogen receptor signaling pathway |
| Ezh2 | CAAAGGATACAGACAGTGACAGAG | CCGAGAATTTGCTTCAGAGGAG | chromatin organization |
| Fgfr2 | CTGCCGCCAACACTGTCAAG | TGACGGGACCACACTTTCCA | animal organ morphogenesis |
| Foxa1 | CACCTTGGTAGTAGGCTGGC | TCCTTATGGCGCTACCTTGC | mes.-epi. cell signaling involved in prostate gland development |
| Hdac1 | AGCCATCTTTAAGCCAGTCATGTC | GAAACTCTTCACGAACTCCACAC | chromatin remodeling |
| Hdac6 | CACCGCATTCAGAGGGTTCT | CCTTAAGGTGGGGCCAGAAG | negative regulation of protein acetylation |
| Hoxa10 | CTCGCTAGTCCCTTTCCTGC | TCTAGGACTCGCTCCTTCC | prostate gland development |
| HoxA7 | GCCTCTGAGGAACCCAGTAGA | CTGGGCCCATAGGTAGTTTGG | multicellular organism development |
| Hoxa9 | AGTTCTCTCCTTGGCGGTTG | AGTCAGACTGGAAAGCCAGC | prostate gland development |
| Hoxb13 | GCCAGATGTGTTGCCAAGGT | GCACAGCCGTCGGGAG | epithelial cell maturation involved in prostate gland development |
| Kat2b | GAGTACCTCTTCACCTGCGT | TGTTCACACCCTGTTCAATACTG | protein acetylation |
| Kiss1 | CGGACCCCAGGAACTCGTTA | GGCATGGCGACGACCTAC | G protein-coupled receptor signaling pathway |
| Kmt2d | CATCCCTGTCTTCCCAGATACCA | CACCTCACTTCCCTTGCCCT | response to estrogen |
| Nkx3-1 | GACCCACCAAGTATCCGGC | CACTTGCTAAGTCCCCTGGATT | prostate gland development |
| Notch1 | CGACAACCGGCAATGTGTGC | CCACCGGCTCACTCTTCACG | prostate gland epithelium morphogenesis |
| Sox9 | AGGCTGTAAAATGCCACTC | CGCTCCGCCTCCTCCACGAA | male gonad development, |
| Srd5a1 | AGGTACCACTGATGATGCTGC | GAGTGGTGTGGCTTTGCACT | androgen biosynthetic process |
| Stat5a | CACGTGGAAGAACTTTTACGCC | AGCATGGAGTCCAGCGTTC | prostate gland epithelium morphogenesis |

### Table S2. Oligonucleotides used for ChIP-qPCR

| gene | forward | reverse | biological process |
|---|---|---|---|
| Akr1b3 | GTTACAAACCCGGACCGGCA | GATCCGTCTCTCAGCCGTGC | enables aldose reductase (NADPH) activity |
| Brca1 | TAAAATTCCCGCGCTCTCCG | CGCTGACGTGTCTGGATCTT | regulation of DNA damage checkpoint |
| Ccnd1 | GGCGGATGGTCTCCACTTC | TCCGGAGACCGGCAGTACA | G1/S transition of mitotic cell cycle |
| Ctnnb1 | CCCAGTAGAGGCCACAGTGC | GACCCAGCAAGGTACAGCCC | Wnt signalling |
| Ep300 | CTCGTGGGATCAGTGTTGCT | GGGATGCGGACTCAACAGAA | animal organ morphogenesis |
| Esr1 | GTTCAACTACCCCGAGGGCG | CCCAGGCTGTTGGCACTGAA | estrogen receptor signaling pathway |
| Ezh2 | TGGTAACGGTCTTAACCGCC | GGTCACACGCCTTCCTTTCA | chromatin organization |
| Fgfr2 | AGTGAGATTCCATCTTCCTCTGGA | CCTCTTCAGCGAAGCGGTTA | animal organ morphogenesis |
| Foxa1 | CCAGCTGATCGGAACCATTCT | GTCACTCCCCGTGGAAAACC | mes.-epi. cell signaling inv. in prostate gland development |
| Hdac1 | GTAAGATGCTCGCGCTGGCT | CATCCCCTCCCCCACTCCAT | chromatin remodeling |
| Hdac6 | CCTCTAATCTGCGCCCGGAAC | GGCGGACTAGAAAGGTGGGTG | negative regulation of protein acetylation |
| Hoxa10 | GCTTCATTACGCTTGCTGCC | CGGTGGAGGTGGCTACTACG | prostate gland development |
| Hoxa7 | GAACCAGGTTTGGGGACCTC | AAGGGAATGAACACCTACGGC | multicellular organism development |
| Hoxa9 | CCGAGAGCGGTTCAGGTTT | CAGGTATATGCGCTCCTGGC | prostate gland development |
| Hoxb13 | TGTGTGCTTTGAGGGAGCCG | GCGCAAGATCAAGCGCAGAC | epithelial cell maturation involved in prostate gland development |
| Kiss1 | CGGACCCCAGGAACTCGTTA | GGCATGGCGACGACCTAC | G protein-coupled receptor signaling pathway |
| Nkx3-1 | CTGCCTGGATCCCGACGTA | AGCTAGCGTCGTAGGGAGTG | prostate gland development |
| Notch1 | TGGTCTCACAGGAGCACCCA | CTTGCCGGGATGGCCTCAAT | prostate gland epithelium morphogenesis |
| Plaur | CCCCGCAGTGAGCGGATAAG | CATCTCGCTGGAGGCTCTGC | epithelial cell differentiation involved in prostate gland |
| Srd5a1 | ACCCTCCAGGTAGACTAGCG | ACTTGGACATCCGAGCATGG | androgen biosynthetic process |
| Stat5a | GGTCATCGATGGAACATGGCTA | TAGGGGATTTGTCATTGTGTGGAT | prostate gland epithelium morphogenesis |

### Table S3. Biological process "prostate gland development"

| | |
|---|---|
| *Ahr* | aryl-hydrocarbon receptor |
| *Apc* | APC, WNT signaling pathway regulator |
| *Ar* | androgen receptor |
| *Bmp4* | bone morphogenetic protein 4 |
| *Cd44* | CD44 antigen |
| *Cdkn1b* | cyclin dependent kinase inhibitor 1B |
| *Ctnnb1* | catenin beta 1 |
| *Cyp19a1* | cytochrome P450, family 19, subfamily a, polypeptide 1 |
| *Cyp7b1* | cytochrome P450, family 7, subfamily b, polypeptide 1 |
| *Eaf2* | ELL associated factor 2 |
| *Esr1* | estrogen receptor 1 (alpha) |
| *Esr2* | estrogen receptor 2 (beta) |
| *Fem1b* | fem 1 homolog b |
| *Fgf10* | fibroblast growth factor 10 |
| *Fgfr2* | fibroblast growth factor receptor 2 |
| *Fkbp4* | FK506 binding protein 4 |
| *Foxa1* | forkhead box A1 |
| *Frs2* | fibroblast growth factor receptor substrate 2 |
| *Gli2* | GLI-Kruppel family member GLI2 |
| *Hoxa7* | zhomeobox A7 |
| *Hoxa9* | homeobox A9 |
| *Hoxa10* | homeobox A10 |
| *Hoxa13* | homeobox A13 |
| *Hoxb13* | homeobox B13 |
| *Hoxd13* | homeobox D13 |
| *Id4* | inhibitor of DNA binding 4 |
| *Igf1* | insulin-like growth factor 1 |
| *Igf1r* | insulin-like growth factor I receptor |
| *Mmp2* | matrix metallopeptidase 2 |
| *Nkx3-1* | NK3 homeobox 1 |
| *Nog* | noggin |
| *Notch1* | notch 1 |
| *Plag1* | pleiomorphic adenoma gene 1 |
| *Plaur* | plasminogen activator, urokinase receptor |
| *Prlr* | prolactin receptor |
| *Psap* | prosaposin |
| *Psapl1* | prosaposin-like 1 |
| *Pten* | phosphatase and tensin homolog |
| *Rarg* | retinoic acid receptor, gamma |
| *Rln1* | relaxin 1 |
| *Rxra* | retinoid X receptor alpha |
| *Serpinb5* | serine (or cysteine) peptidase inhibitor, clade B, member 5 |
| *Serpinf1* | serine (or cysteine) peptidase inhibitor, clade F, member 1 |
| *Sfrp1* | secreted frizzled-related protein 1 |
| *Shh* | sonic hedgehog |
| *Sox9* | SRY (sex determining region Y)-box 9 |
| *Stat5a* | signal transducer and activator of transcription 5A |
| *Stk11* | serine/threonine kinase 11 |
| *Tnc* | tenascin C |
| *Trp63* | transformation related protein 63 |
| *Ube3a* | ubiquitin protein ligase E3A |
| *Wdr77* | WD repeat domain 77 |
